# Regularly updated benchmark sets for statistically correct evaluations of AlphaFold applications

**DOI:** 10.1101/2024.08.02.606297

**Authors:** Laszlo Dobson, Gábor E. Tusnády, Peter Tompa

**Affiliations:** Institute of Molecular Life Sciences, Research Centre for Natural Sciences, Magyar Tudósok Körútja 2, Budapest, 1117, Hungary; Department of Bioinformatics, Semmelweis University, Tűzoltó u. 7, Budapest, 1094, Hungary; VIB-VUB Center for Structural Biology, Vlaams Instituut voor Biotechnologie (VIB), 1050 Brussels, Belgium; Structural Biology Brussels (SBB), Vrije Universiteit Brussel (VUB), 1050 Brussels, Belgium

**Keywords:** alphafold, benchmark set, data leakage, disordered proteins, structure prediction

## Abstract

AlphaFold2 changed structural biology by providing high-quality structure predictions for all possible proteins. Since its inception, a plethora of applications were built on AlphaFold2, expediting discoveries in virtually all areas related to protein science. In many cases, however, optimism seems to have made scientists forget about data leakage, a serious issue that needs to be addressed when evaluating machine learning methods. Here we provide a rigorous benchmark set that can be used in a broad range of applications built around AlphaFold2/3.

**Graphical abstract:** 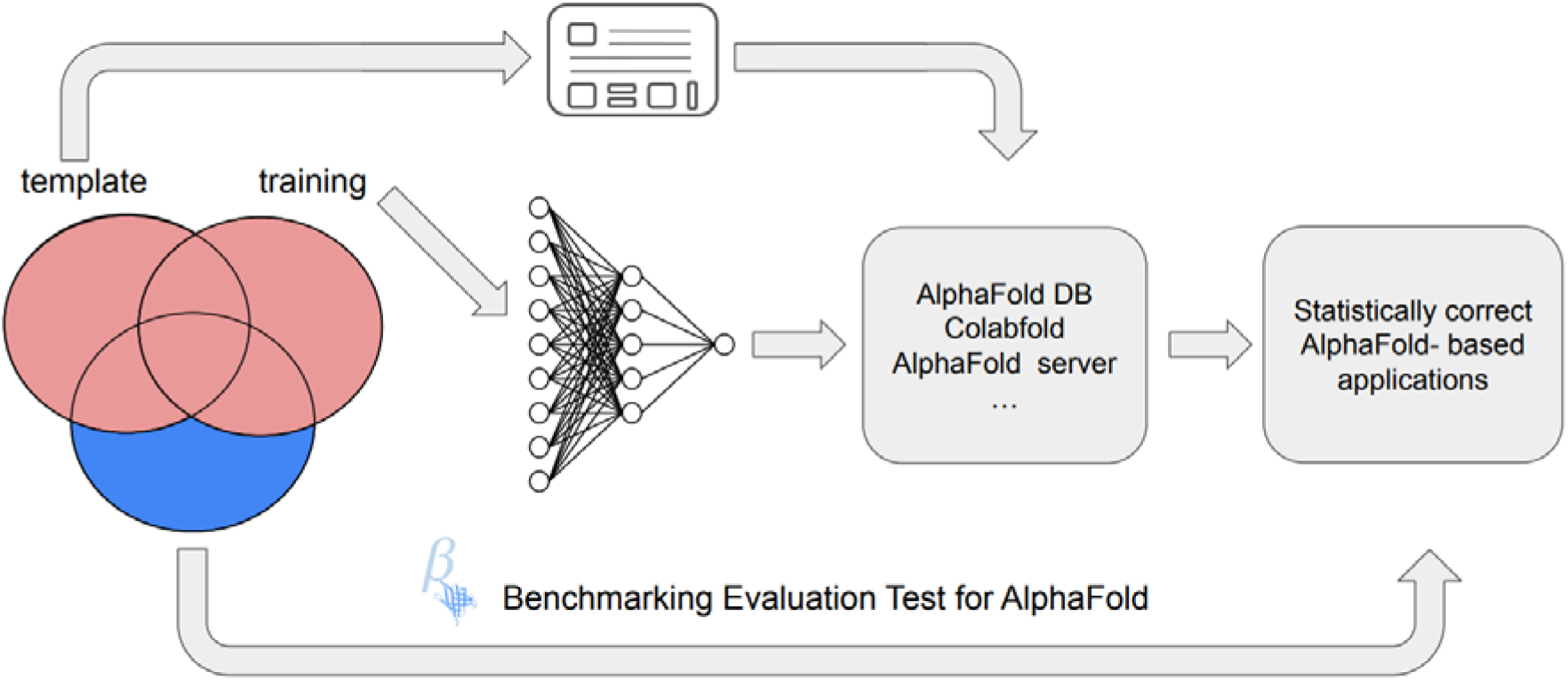

*Key Points:*

- When building applications on AlphaFold, scientists should consider the possibility of data leakage between AlphaFold training set and the independent test set of their method
- BETA provides multiple datasets with structures and sequences that were not used during the training of AlphaFold
- These datasets provide a diverse range of use cases
- The protocol was applied when building a simple disordered prediction method, showing different parameters required to optimize disordered prediction for proteins not used in AlphaFold training

## Introduction

The creation of AlphaFold2 (AF2) in 2020 was bound to revolutionize protein science, by significantly raising the standards of structure prediction at CASP XIV [1]. Within a year, the method was published [2], soon followed by a database of predicted all-atom structures for almost every known protein sequence [3]. Based on this spectacular feat, thousands of papers appeared that analyzed AF2 structures to predict and/or assess various structural and physicochemical features of proteins in specific areas, such as protein stability [4], conformational heterogeneity [5], protein function [6], structure [7] or interfaces [8] of protein complexes, domain definitions [9], or disordered regions that do not fold into unique structures [10]. A similar, or even more sweeping campaign of applications can be expected now with the appearance of AF3, with its advanced architecture and features, probably more reliably predicting protein-ligand (another protein), protein-nucleic acid and protein-small molecule interactions and complex structures [11].

The apparent insight provided by these noted, and many other original applications gave cause for optimism, having reinvigorated a broad range of structure-related studies, with artificial intelligence promising to expand the proteome coverage enabled by laborious and time-consuming experiments. We noted, however, that many of these studies did not take a stringent approach to selecting proteins that had not been used to train AF2, leading to data leakage, which is inevitably leads to statistical inaccuracies when applying machine learning methods. Most of these papers are still highly valuable and point to relevant applications and evaluations. It is of relevance that our perusal of literature could not be done right after AF2 publication, only after an intense period of diverse analyses based on its applications. We figure the same will happen after AF3 code and/or access will be made broadly available, thus our conclusion and suggested solution also apply across a wide range of future applications.

AF2 has not been trained on protein complexes, its performance in predicting their structures is remarkable, probably because protein-protein interactions have a similar biophysical background to protein folding, which AF2 has been trained to model – these capabilities are promised to further improve with AF3. Intriguingly, the capabilities of AF2 to model complexes is not limited to those built on domain-domain interactions [7], it can also well handle more transient interactions driven by short linear motifs [9]. It should be noted, however, that although the evaluation of multimers appears to be not affected by the choice of benchmarks, the bound state of the respective single chains used for the training of AF2 does cause data leakage and compromises the resulting statistical inference.

To overcome this limitation we suggest a deep homology search for proteins in the training/template sets of AF. Although the solution may seem trivial, considering how distant homologous structures can be still captured by AF and the many different versions of AF each with a different training cutoff, we prepared benchmark sets to facilitate this process. Here we provide Benchmarking Evaluation Test for AlphaFold (BETA), containing predefined datasets for AF applications, and demonstrate through a range of case studies that it resolves the issue of data leakage-derived statistical uncertainties in AF-based analyses.

## Methods

### Defining benchmark sets

BETA was created as follows. PDB [6], SwissProt [12] and BIOGRID [13] were downloaded on the 21th of May, 2024. Considering the first day of the following month of each cutoff date (30.04.2018; 31.05.2020; 15.02.2021; 30.09.2021; 15.07.2022, 01.11.2022, 01.01.2023, 01.01.2024) we performed homology searches based on both sequence and structure. AF can be also used without incorporating template structures, and for training AF NMR structures were not used as models, therefore for cutoff dates related to training (30.04.2018,30.09.2021) NMR structures were not included in the background database. These dates were selected based on how different AF versions training/template libraries were selected (see Figure 1). Homology searches were carried out as follows: I) Sequence search (PSI-BLAST [14]) of all structures after cutoff date against structures before cutoff date; II) Structure search (Foldseek [15]) of all structures after the cutoff date against structures before cutoff date; III) Sequence search (PSI-BLAST) of SwissProt proteins against structures before cutoff date. In the case of PSI-BLAST (e-value 0.0001, 3 iterations, maximum target sequences 50 000), all queries having any hit longer than 10 amino acids with at least 20% sequence identity were filtered In Foldseek searches (maximum target structures 50 000), all queries having any hit longer than 10 amino acids with at least 0.25 TM-Score were filtered. We used Voronota [16] to detect all interchain interaction in PDB structures, considering the most probable oligomerization state stored on PDBe. We used BIOGRID (only ‘direct interactions’) to search for interactions in SwissProt sequences. The pipeline and relevant resulting collections of different BETA benchmark sets are shown on Figure 1.

**Figure 1:**
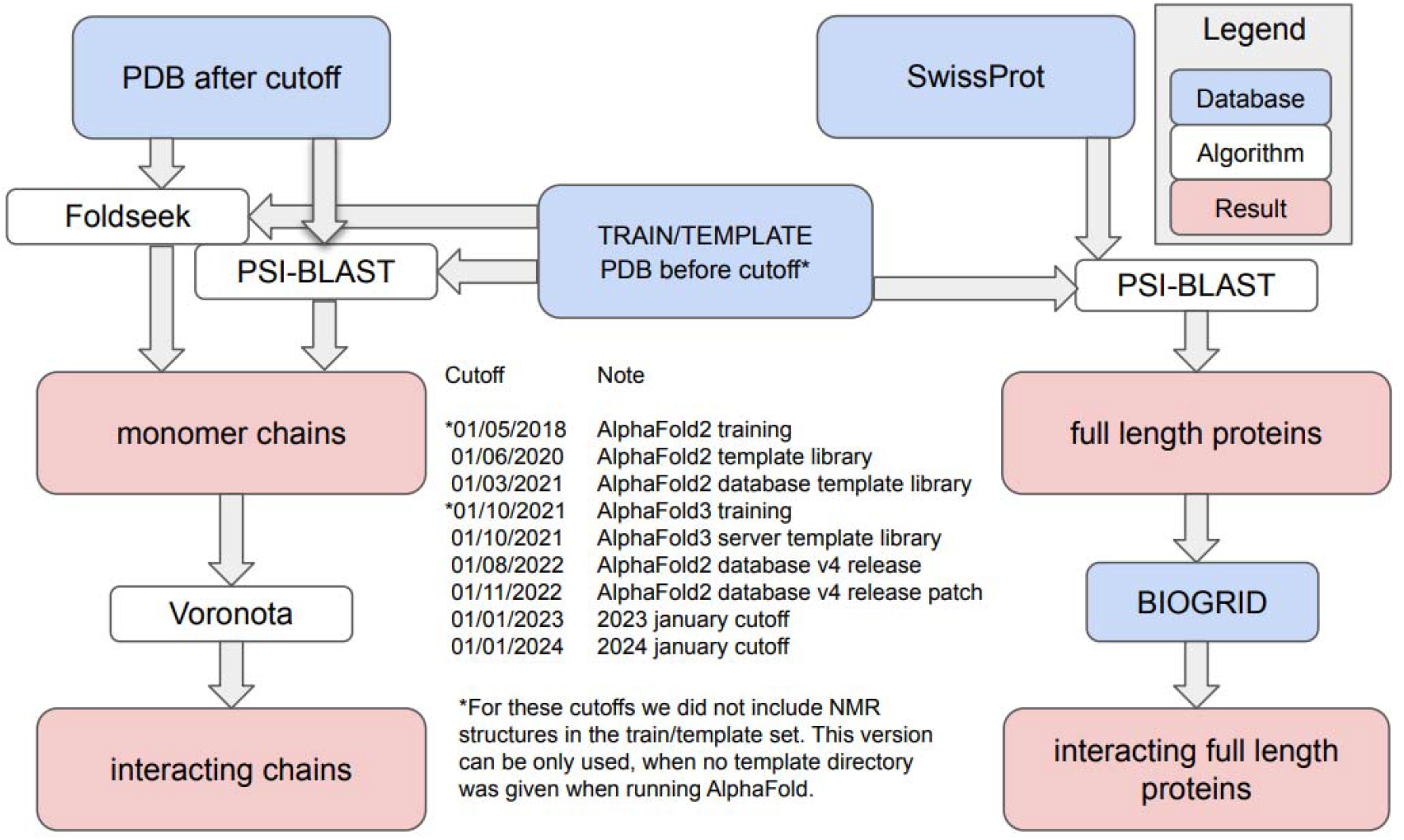
Pipeline used to define the Benchmarking Evaluation Test for AlphaFold (BETA). Blue and white panels represent databases and algorithms (respectively), red panel show results of the pipeline. Predefined cutoff dates and their rationale can be found in the middle.

### A case study: prediction of protein disorder

To demonstrate the importance of our methodology, we assessed how prediction methods of intrinsically disordered proteins/regions (IDPs/IDRs) would be affected by utilizing BETA (Figure 2/A). Previously, the confidence measure Local-Distance Difference Test (pLDDT) was shown to be a good predictor of protein disorder [10]. We downloaded all PDB structures, where the most probable oligomeric state was a monomer according to PDBe [6]. Sequences were filtered to 40% identity using CD-HIT [17], and disorder of residues was inferred, if coordinates of their side-chains were missing from the PDB structures. By adhering to the straightforward definition of missing coordinates, we: i) accept the consensus in the IDP community, as reflected in state-of-the-art protein disorder annotation databases DisProt [18] and MobiDB [19]; ii) define a straightforward and clear methodology, so readers/users do not get lost in details of the pipeline; and iii) use an approach where no cutoffs required (that can be always argued). As the resolution of structures may have no effect on the choice of disordered residues [20], at the moment it is not considered in standard approaches of selecting for disordered residues, for example, in the latest release of the DisProt database [18] or in formulating respective community guidelines [21]. Also, in accord with guidelines, segments shorter than 10 amino acids were omitted, and the remaining residues (with side-chain coordinates) were labeled as ordered. We used SIFTS [22] to cross-reference UniProt identifiers and positions with PDB chains and residues, then downloaded AF2 structures from the AlphaFold protein structure database. Structures were categorized into ‘Homologous structures’ and ‘BETA structures’, considering the AF database training/template cutoff, which in this case is 01/03/2021. Supplementary Table 1 contains all residues incorporated in the analysis (with cross-referenced PDB and AF positions), with ordered and disordered residue classifications (inferred from PDB structures), AF2 pLDDT values, as well as BETA and homologous structure classifications.

**Figure 2.**
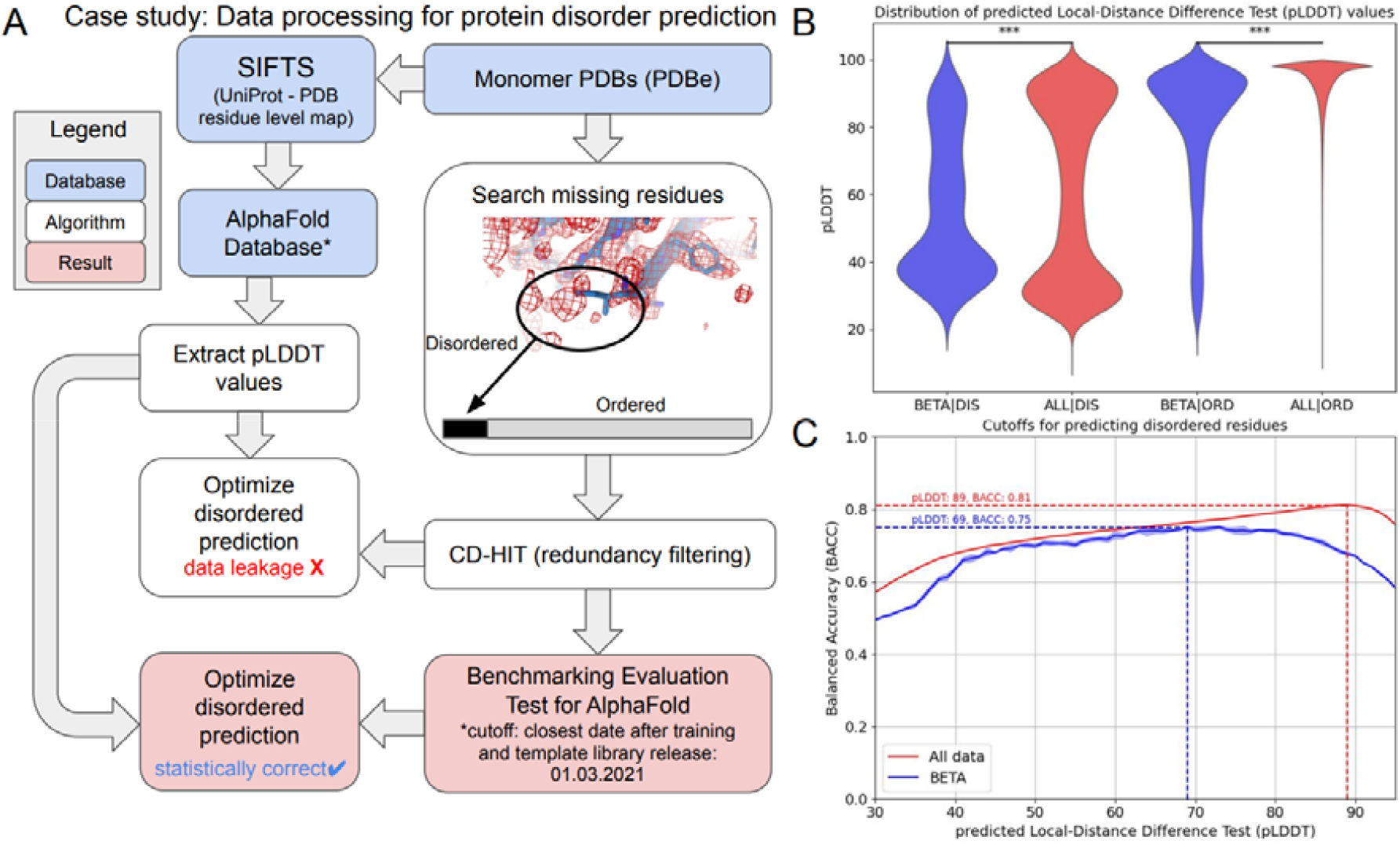
Pipeline and evaluation of the protein disorder prediction case study. A) Pipeline to define disordered residues from PDB and incorporating BETA into the pipeline. For more details, see Methods. B) Distribution of pLDDT values of disordered and ordered residues in all monomeric structures (red) and BETA structures (blue). There is highly significant difference between pLDDT values in the case of BETA against all structures, both for ordered and disordered residues. C) Correspondence between pLDDT and Balanced accuracy considering all structures and BETA structures. The highest Balanced accuracy (with suggested pLDDT cutoff is shown). The small area around curves show the standard error calculated from bootstrapping 50% of the data 5 times, while the curves show the mean value at a given point.

## Results

### Benchmarking Evaluation Test for AlphaFold (BETA)

To place AF2 – and by the same logic, AF3 – based structure-function studies on a solid ground, we present here the Benchmarking Evaluation Test for AlphaFold (BETA), a dataset of structures and sequences that were not used in AF training (https://beta.pbrg.hu, https://github.com/brgenzim/Beta). In BETA, we list (i) monomeric PDB chains with no homologs before the selected date; (ii) interacting PDB chain pairs where participants have no homologs prior the selected date; (iii) complete protein sequences from SwissProt without any homologs in the PDB before the selected date and (iv) complete interacting sequences from SwissProt where the interacting participants do not have homologs before the selected date (Figure 1).

BETA can be utilized in many different ways to help construct a benchmark set for surveys or prediction methods, while also providing a convenient way for researchers without a computational background to browse data. We aimed to construct datasets for all users, including those accessing AF via the AlpaFold database [3], ColabFold [23] or the AlphaFold server [11], or running their predictions on their own computer. For defining an appropriate benchmark set using BETA, one should first determine the appropriate cutoff date based on the input data (UniProt sequences or PDB codes) and the version of AF (AlphaFold database, ColabFold, AlphaFold server or running on one’s own computer), and then select the first date after this cutoff to avoid using structures used during AF2/AF3 training or prediction. To demonstrate the general applicability of BETA, we outline a highly informative case study of predicting structural disorder, out of five relevant Use Cases (more details are available at Supplementary Material S1 and https://beta.pbrg.hu/): (1) Building a protein disorder prediction method (further detailed as a case study); (2) Predicting antibody epitopes; (3) Predicting phase-separating regions in proteins; (4) Assessing the pathological effects of missense mutations; and (5) Ranking interactions driven by short linear motifs. These examples explain how an appropriate benchmark dataset can be selected and used.

### Case study: the effect of using BETA for protein disorder prediction

To set the precise pLDDT threshold to predict IDRs, we can define a set of proteins where we can assign labels to each residue, whether they belong to IDRs or not. Using a benchmark set of PDB structures (see Methods), we looked for the distribution of pLDDT values in ordered and disordered residues (Figure 2/B, red distributions). As the two distributions are different, one can assume that pLLDT value is an indeed good predictor for protein disorder. However, limiting the examined residues to those that are available in the BETA database, there is also a significance difference between ordered residues in BETA and all ordered residues (Kolmogorov-Smirnov (KS) Test, p<0.0001), as well as between residues in IDRs (BETA against all structures, K-S Test, p<0.0001) (Figure 2/B). To further highlight the gravity of differences caused, we also looked for the optimal pLDDT cutoff to achieve the highest balanced accuracy (Figure 2/C, Supplementary Table 2). Indeed, the cutoff for the highest accuracy is different for all structures (pLDDT=0.89) from the cutoff defined using only BETA structures (pLDDT=0.69). Notably, the number of residues in IDRs is relatively low in the BETA set defined by monomeric PDBs (total 1062 residues), which is likely caused by the fact that most novel structures deposited to PDB correspond to big complexes. To avoid statistical bias, we randomly sampled 50% of the residues five times to define the pLDDT cutoffs.

We have to add that we do not aim to build a perfect IDR predictor here, and these steps can be further optimized. Our goal was to show that defining a stringent benchmark set may heavily influence an application built on AF, both in the selected cutoff, but also on the measured accuracy which is lower when data leakage is ruled out.

## Conclusion

While AF2 is already available through multiple resources, AF3 is currently available as a web server (or the standalone program), with a broad range of applications for predicting various features on the horizon. We noticed that many applications built on AF do not take into account the training set of AF. Our comparison on predicting protein disorder demonstrates that our conclusion and suggestion are useful for future applications of AF, thus we plan to update the underlying BETA dataset on a regular basis, taking into account how AF2/AF3 based resources will update their training and template libraries in the future. Of possibly even broader relevance of applicability, multiple other structure prediction methods have been published since AF was developed [24-26]. Although the website of BETA cannot offer an exact cutoff date for the training date of each method, researchers can still rely on the website (with the nearest training cutoff) or perform homology search with custom cutoff dates using the GitHub repository, when building applications on structure prediction tools.

## Authors

Laszlo Dobson is a researcher at the HUN-REN Research Centre for Natural Sciences. His research involves computational biology and the study of protein-protein interactions and membrane proteins.

Gabor Tusnady is a group leader at the HUN-REN Research Centre for Natural Sciences. His research area are investigation of membrane protein topology and 3D structure by developing computational algorithms and maintaining databases and servers for transmembrane protein structure and topology.

Peter Tompa is a professor at the VIB-VUB Center for Structural Biology. His work focuses on intrinsically disordered proteins and phase-separation.

## Data availability

Supplementary Table 1 is available at https://zenodo.org/records/14711867. Supplementary Table 2 is available Online.

## Acknowledgment

We thank Rita Pancsa for discussing the manuscript and Zsofia Kalman for the website logo.

