## Supplementary material for "Regularly updated benchmark sets for statistically correct evaluations of AlphaFold applications": Supplemantary Material S1

The following five use cases describe how to define a benchmark set using BETA.

**Use case 1: Building a protein disorder prediction method**
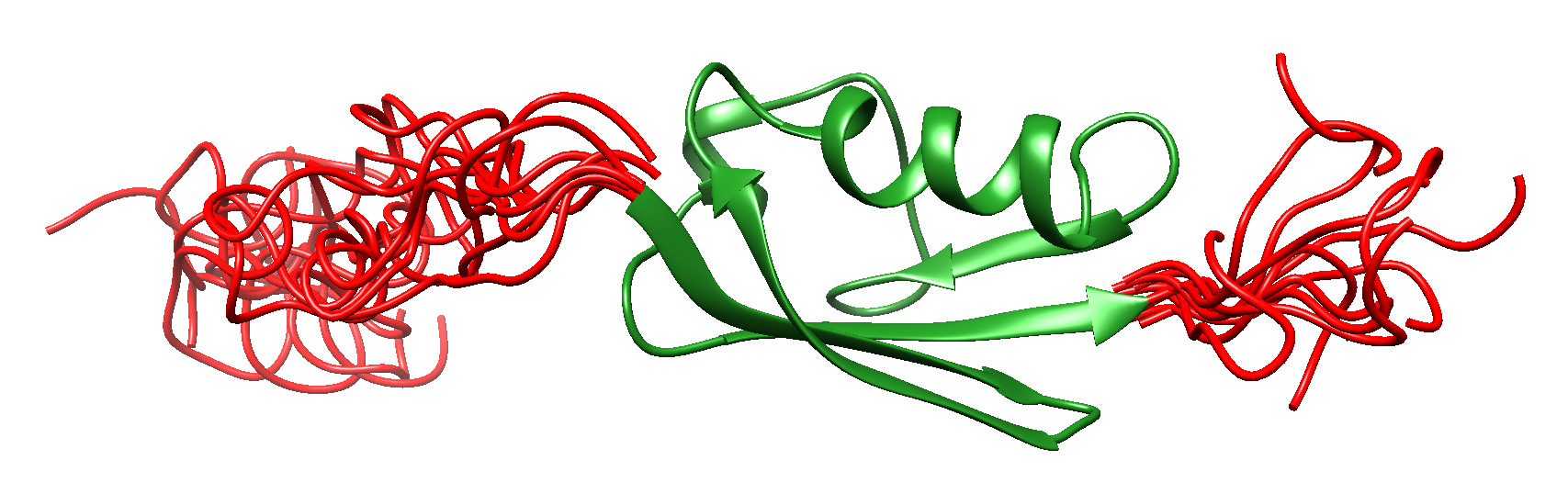

**Problem:** We would like to build a protein disorder prediction method based on AlphaFold2 structures, utilizing accessibility and pLDDT value of residues. To fine tune parameters we prepared a dataset of disordered regions from DisProt [<https://disprot.org/>] and also prepared a dataset of ordered monomeric proteins from PDB [<https://www.ebi.ac.uk/pdbe/>] that we mapped to UniProt [https://www.uniprot.org/], using SIFTS [<https://www.ebi.ac.uk/pdbe/docs/sifts/>]. We don’t have the capacity to run AlphaFold2 on our computers, so we use AlphaFold Database [<https://alphafold.ebi.ac.uk/>] to download precomputed AlphaFold2 structures.

**Initial dataset:** We have a list of UniProt ACs containing disordered and ordered regions/proteins.

**Protocol to follow:**

1) AlphaFold database [<https://alphafold.ebi.ac.uk/>] uses AlphaFold version trained [<https://alphafold.ebi.ac.uk/faq>] on structures released before 30.04.2018, but also utilizes template structure before 15.02.2015.

2) Go to BETA download page [<https://beta.pbrg.hu/download>] and select the benchmark version. We have UniProt AC lists, so for the benchmark set we need full length proteins. For the cutoff date we select the closest date after training and template library release, that is 01.03.2021.
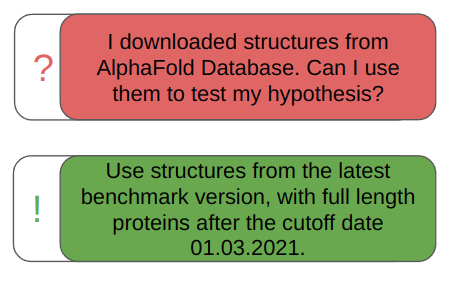

3) The intersection of the original benchmark set (steps 1) and the filtered set from BETA (step 2) should be used to fine tune cutoff values.

**Use case 2: Predicting antibody epitopes**

**Problem:** We would like to assess whether AlphaFold3 can predict antibody epitopes. We use SabDab [<https://opig.stats.ox.ac.uk/webapps/sabdab-sabpred/sabdab>] to collect structures. We use AlphaFold3 server [<https://golgi.sandbox.google.com/>] to predict the structures.
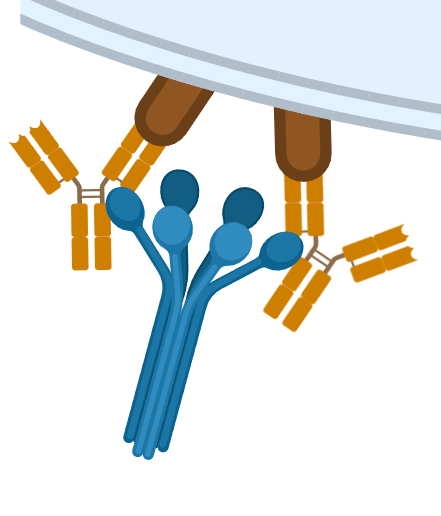

**Initial dataset:** We have a list of PDB chain pairs containing antibody-antigene interactions.

**Protocol to follow:**

1) According to the FAQ provided to AlphaFold3 [<https://golgi.sandbox.google.com/faq>] the server does not use structures submitted to PDB after 30.09.2021.

2) Go to the BETA webpage [https://beta.pbrg.hu/] On the navbar above type your PDB code into the search input field, then select the closest date after training and template library release, that is 01.10.2021.
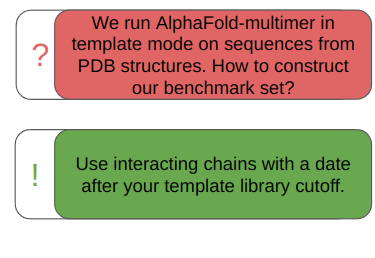

3) The intersection of the original benchmark set (step 1) and the filtered set from BETA (step 2) should be used to evaluate predictions.

**Use case 3: Preciting phase-separating regions in proteins**

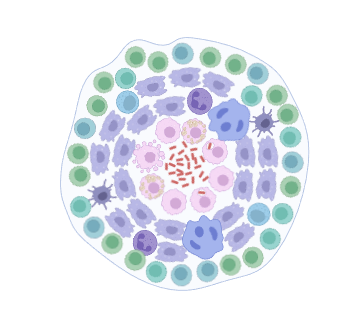

**Problem:** We have a list of a few proteins undergoing self-associated phase-separation. We would like to see whether AlphaFold2 can highlight relevant - predictive – features of regions of these proteins. We don’t have a computational background, we just want to objectively judge a handful of structures from AlphaFold Database [<https://alphafold.ebi.ac.uk/>]. We managed to collect UniProt [<https://www.uniprot.org/>] ACs for our proteins.

**Initial dataset:** We have a list of UniProt ACs containing disordered and ordered regions/proteins.

**Protocol to follow:**

1) AlphaFold database [<https://alphafold.ebi.ac.uk/>] uses AlphaFold version trained [<https://alphafold.ebi.ac.uk/faq>] on structures released before 30.04.2018, but also utilizes template structure before 15.02.2021.
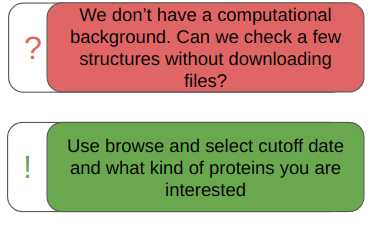

2) Go to the BETA browse page [<https://beta.pbrg.hu/browse>] and type your UniProt AC, then select the closest date after training and template library release, that is 01.03.2021.

3) If your structure can be used for evaluation, a green checkmark will appear.

**Use case 4: Assessing the pathological effects of missense mutations**

**Problem:** We would like to use AlphaFold2 to decide on the possible pathological outcome of missense variations. We have a list of mutations with outcome from UniProt humsavar [<https://ftp.uniprot.org/pub/databases/uniprot/current_release/knowledgebase/complete/docs/humsavar.txt>]. We download AlphaFold [<https://github.com/google-deepmind/alphafold>] and predict wild type and variant structures on our own computer, without using templates. Then we use other tools to assess stability of the predicted structures.

**Initial dataset:** We have a list of UniProt ACs from humsavar together with information about variants.
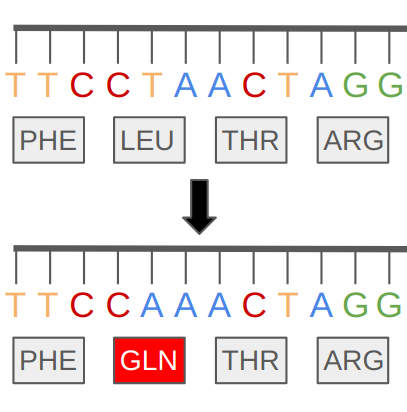

**Protocol to follow:**

1) The original AlphaFold was trained on structures deposited to PDB before 30.04.2018 [<https://www.nature.com/articles/s41586-021-03819-2>]. We don’t use templates.

2) Go to the BETA download page [<https://beta.pbrg.hu/download>] and select benchmark version. For the benchmark set we use full length proteins. For date we select the closest date after the cutoff, which is 01.05.2018.
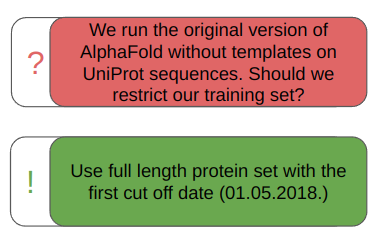

3) The intersection of the original benchmark set (steps 1) and the filtered set from BETA (step 2) should be used to calculate stabilities.

**Use case 5: Ranking interactions driven by short linear motifs**

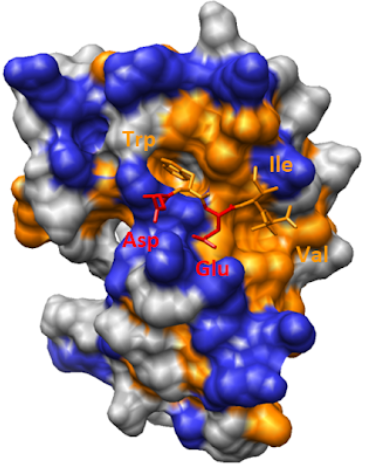

**Problem:** We would like to predict whether AlphaFold2 scores can be used to rank short linear motif based interactions. We have an experimental setup that use different peptides as bait and use protein domains as prey. We would like to compare assay results with computational predictions. Using BLAST and peptide search we prepared a list of UniProt [<https://www.uniprot.org/>] ACs, containing domains and peptides. We use Colabfold [<https://colab.research.google.com/github/sokrypton/ColabFold/blob/main/AlphaFold2.ipynb>] with default settings to predict combinations of different interactions. We do not use template structures.

**Initial dataset:** We have a list of UniProt identifiers, some of them were shown to bind to each other.

**Protocol to follow:**

1) ColabFold with default settings uses AlphaFold2-Multimer. According to the README provided to AlphaFold [<https://github.com/google-deepmind/alphafold?tab=readme-ov-file>] the model for the multimer version was retrained on 30.09.2021.
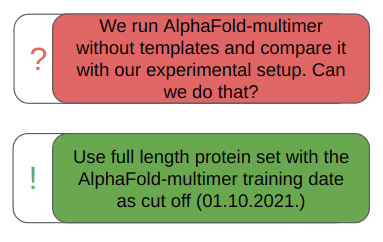

2) Go to the BETA download page [<https://beta.pbrg.hu/download>] and select the benchmark version. For the benchmark set we use full length proteins. For date we select the closest date after the cutoff, which is 01.10.2021.

3) The intersection of the original benchmark set (step 1) and the filtered set from BETA (step 2) should be used to assess predictions.

*(4) Alternatively, interacting full length proteins can be also selected to compare results with interactions stored in BIOGRID*
